# Synchronous Measurements of Extracellular Action Potentials and Neurochemical Activity with Carbon Fiber Electrodes in Nonhuman Primates

**DOI:** 10.1101/2023.12.23.573130

**Authors:** Usamma Amjad, Jiwon Choi, Daniel J. Gibson, Raymond Murray, Ann M. Graybiel, Helen N. Schwerdt

## Abstract

Measuring the dynamic relationship between neuromodulators, such as dopamine, and neuronal action potentials is imperative to understand how these fundamental modes of neural signaling interact to mediate behavior. Here, we developed methods to measure concurrently dopamine and extracellular action potentials (i.e., spikes) and applied these in a monkey performing a behavioral task. Standard fast-scan cyclic voltammetric (FSCV) electrochemical (EChem) and electrophysiological (EPhys) recording systems are combined and used to collect spike and dopamine signals, respectively, from an array of carbon fiber (CF) sensors implanted in the monkey striatum. FSCV requires the application of small voltages at the implanted sensors to measure redox currents generated from target molecules, such as dopamine. These applied voltages create artifacts at neighboring EPhys-measurement sensors, producing signals that may falsely be classified as physiological spikes. Therefore, simple automated temporal interpolation algorithms were designed to remove these artifacts and enable accurate spike extraction. We validated these methods using simulated artifacts and demonstrated an average spike recovery rate of 84.5%. This spike extraction was performed on data collected from concurrent EChem and EPhys recordings made in a task-performing monkey to discriminate cell-type specific striatal units. These identified units were shown to correlate to specific behavioral task parameters related to reward size and eye-movement direction. Synchronous measures of spike and dopamine signals displayed contrasting relations to the behavioral task parameters, as taken from our small set of representative data, suggesting a complex relationship between these two modes of neural signaling. Future application of our methods will help advance our understanding of the interactions between neuromodulator signaling and neuronal activity, to elucidate more detailed mechanisms of neural circuitry and plasticity mediating behaviors in health and in disease.

**Significance statement:** We present a simple method for recording synchronous molecular and neuronal spike signals. Conventional electrophysiological and electrochemical instruments are combined without the need for additional hardware. A custom-designed algorithm was made and validated for extracting neuronal action potential signals with high fidelity. We were able to compute cell-type specific spike activity along with molecular dopamine signals related to reward and movement behaviors from measurements made in the monkey striatum. Such combined measurements of neurochemical and extracellular action potentials may help pave the way to elucidating mechanisms of plasticity, and how neuromodulators and neurons are orchestrated to mediate behavior.

## Introduction

Most neurons communicate with each other via both chemical and electrical forms of signaling. The relative timing between dopamine neurotransmitter release and action potentials of target neurons has been shown to regulate synaptic plasticity (Brzosko et al., 2019; Reynolds et al., 2001; Shindou et al., 2019; Yagishita et al., 2014). Thus, measuring how chemical and electrical neural signals are coordinated during online behavior is central to studying brain function and the multi-modal mechanisms that shape how neural circuits are programmed to regulate behavior.

Nevertheless, most of our understanding of brain function comes from measurements of either chemical or electrical signals, but not both. Such single mode measurements preclude the ability to look at how molecules, such as dopamine neurotransmitters, modulate nearby neuronal spike activity, and vice-versa—how neuronal activity influences extracellular molecular signaling. Furthermore, there is a critical gap in our knowledge of how such signals are coordinated to mediate behavior. Thus, dual electrical and chemical measurements are imperative to dissect these interactions, especially in awake behaving animals.

Such dual recording may be achieved by combining standard electrophysiology (EPhys) with molecular recording techniques. Molecular measurements may involve electrochemical or optical methods that provide the spatiotemporal resolution to capture the millisecond dynamics of neurotransmitter release and clearance within microscale domains (Rice et al., 2011). Optical methods have been introduced recently, and advanced immensely in use over the past few years (Patriarchi et al., 2018; Sun et al., 2018). These methods require genetic modification of neurons to express synthetic fluorescent receptors (e.g., dLight and GRABDA), and/or introduction of exogenous compounds (e.g., fluorescent false neurotransmitters) (Gubernator et al., 2009; Pereira et al., 2016) or cells (e.g., CNiFER) (Nguyen et al., 2010). Fluorescent approaches demonstrate remarkable specificity for a variety of neurotransmitters and neuromodulators such as dopamine, serotonin, acetylcholine, and many more compounds (Jing et al., 2020; Unger et al., 2020). Nevertheless, application in nonhuman primates has been limited, and such genetic modification is prohibited in humans due to ethical reasons.

This work uses electrochemical (EChem) methods, specifically, fast-scan cyclic voltammetry (FSCV), which has been established over several decades (Clark et al., 2010; Kawagoe et al., 1993) to record current generated through reduction and oxidation (i.e., redox) of dopamine and other chemical compounds. Voltage is scanned at the implanted electrode to generate this redox current at analyte-specific voltages, and this current is proportional to the concentration of the analyte. Carbon fiber (CF) electrodes (CFEs) have been the mainstay for such sensors given their high electron transfer, biocompatibility, and adsorptive properties (McCreery, 2008). Other innovative materials and structures have also shown great potential for enhancing measurement sensitivity and selectivity (Cuniberto et al., 2021; Schmidt et al., 2013). Chemical specificity is determined by the redox voltages, and these voltages are also dependent on the sensor material and scan parameters. Parameters optimal for selective and sensitive dopamine detection have been established over several decades (Keithley and Wightman, 2011; Rodeberg et al., 2017). These electrochemical techniques are also capable of providing readouts of multiple chemical entities by virtue of each compound’s distinct redox voltages (Calhoun et al., 2018; E. Dunham and Jill Venton, 2020; Nguyen and Venton, 2014).

However, use of standard scan parameters (400 V/s) and materials (CF) does not allow distinction of molecules with similar chemical structures (e.g., dopamine and norepinephrine). Another limitation of FSCV is the inability to directly record non-electroactive compounds (e.g., acetylcholine or glutamate). On the other hand, there are ways to indirectly measure these compounds using enzyme coatings that convert these to current-generating H_2_O_2_ (Asri et al., 2016; Kimble et al., 2023). Finally, sensor “fouling” has been a challenge as it relates to the sensitivity degradation of the implanted sensor, and is caused by the semi-permanent adsorption of redox-reaction byproducts or other molecules in the parenchyma (but, see (E. Dunham and Jill Venton, 2020) for methods to address such issues). Nevertheless, FSCV has demonstrated robust performance for chronic measurements in rodents (Clark et al., 2010; Schwerdt et al., 2018) and monkeys (Schwerdt et al., 2020, 2017), and has even been applied in humans intra-operatively (Kishida et al., 2016). Thus, the realization of its potential for clinical translation is imminent.

Concurrent measurement of molecular and electrical neuronal activity is performed by combining EChem and EPhys, respectively, in this work, and will be referred to hereon as ECP (i.e., electrochemical and electrophysiological) recording for simplicity (**Fig. 1**). ECP measurements are not straightforward due to the interference caused by FSCV (**Fig. 1B**). FSCV operates by applying a triangular time-varying voltage (–0.4 to 1.3 V) to the implanted sensor to induce voltage-dependent redox of molecules at its surface; This is usually done at a sampling rate of 10 Hz (i.e., scans are applied every 100 ms). These applied voltages may be directly picked up by the EPhys system in the form of voltage artifacts. Removing these artifacts from the analysis is imperative to compute accurate local field potential (LFP) and spike information in the EPhys recording. Scan artifacts can be erroneously labeled as spikes using standard thresholding and clustering-based spike sorting methods (Schmitzer-Torbert et al., 2005; Schmitzer-Torbert and Redish, 2004). Consequent analytical measures, like spike firing rates, are then susceptible to error and misrepresentation of the actual neural environment. The level of interference depends on how these two measurements are combined; ECP may be performed on the same sensor, or on separate sensors inside the brain.

**Figure 1.**
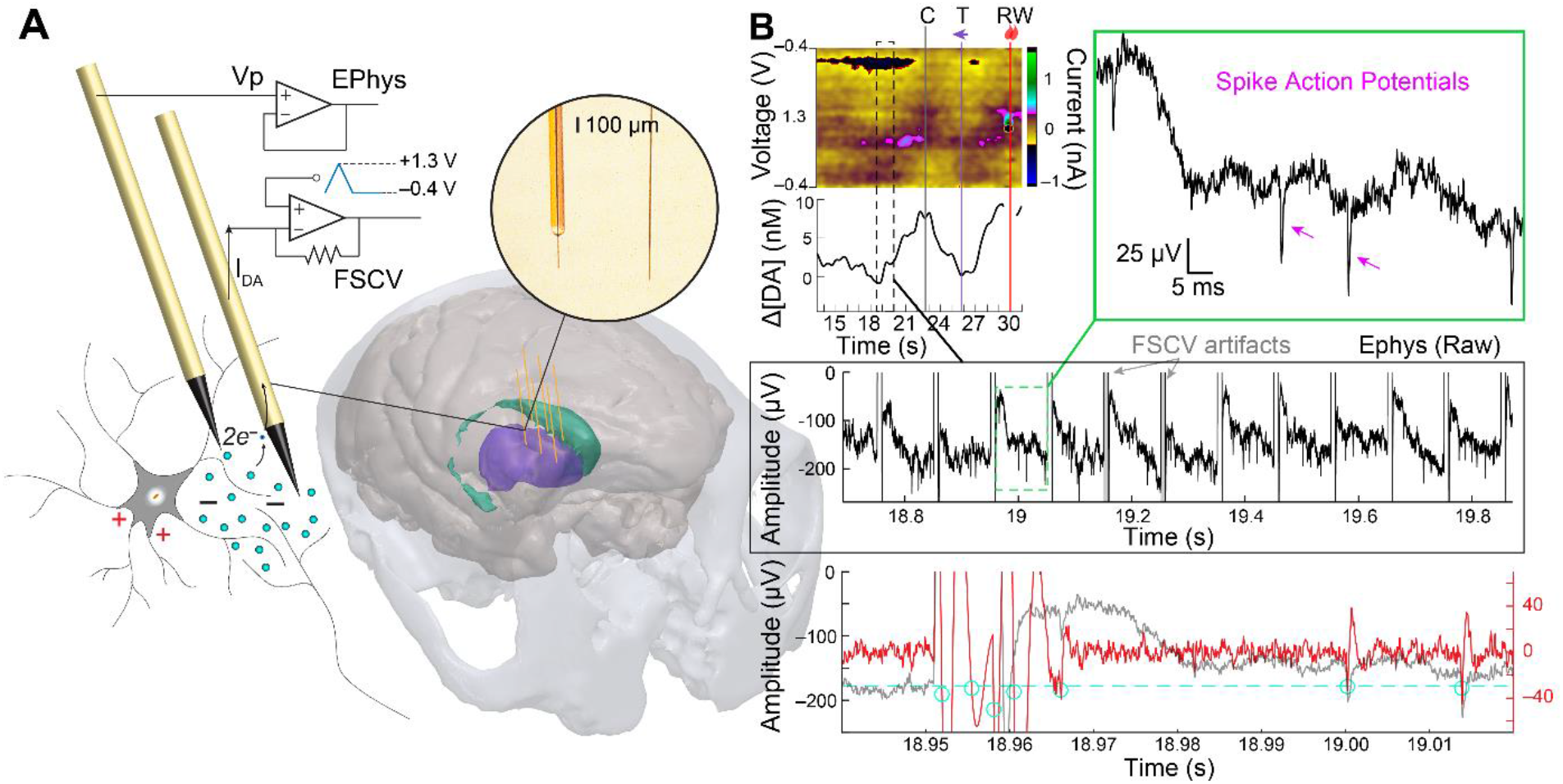
(A) ECP recording setup for synchronous recording of dopamine and neuronal action potentials as recorded from separate CF electrodes implanted in the monkey striatum (colored purple and green) and connected to electrochemical (FSCV) and EPhys recording systems, respectively. (B) Example recordings from ECP system in a task-performing monkey. FSCV-recorded dopamine signals are plotted as a function of time as displayed on a color plot where current changes (color scale) are clearly observed at the redox potentials for dopamine (∼–0.2 V and 0.6 V) and its PCA-extracted dopamine concentration change ([ΔDA]) below it. Task events for the display of the initial central cue (C), peripheral reward-predictive target (T), and reward delivery (RW) are displayed as vertical lines. Below this, the concurrent EPhys recording is shown for a small time window (black dashed rectangle) during the dopamine trace, showing the interfering FSCV scan artifacts. A close-up of the EPhys recording between two scan artifacts (green dashed rectangle) is shown to visualize clear spike action potentials (arrowheads on top right inset). The bottom panel shows the signal after high-pass filtering at 250 Hz using forward-only filters as would be applied for standard spike detection algorithms. This period includes a close-up of the scan artifact demonstrating its triggering of multiple threshold (dashed line)-crossings (circles). Three physiological units are also detected, but the first of these is largely distorted by the forward-filtering of the artifact.

Recording EPhys and FSCV simultaneously from the same sensor usually requires actively switching between the recording modalities so that the applied voltage from FSCV does not overly saturate and/or damage the input amplifiers on the EPhys system. This temporal multiplexing is possible because standard FSCV parameters optimized for dopamine detection use only an 8.5 ms period of the 100 ms sampling interval for applying the redox-generating voltage scan; the remaining 91.5 ms interval is used to hold a negative potential (e.g., –0.4 V) to attract positively charged molecules such as dopamine to the CF (Bath et al., 2000). However, applying a holding potential would effectively short the EPhys input in between scans when applied at the same sensor.

Instead, FSCV and EPhys may be applied at separate sensors; this decoupled ECP configuration permits applying a holding potential, and, in addition, may remove the need for supplementary circuits to switch between EPhys and FSCV operations since the applied voltage is attenuated as a function of distance to other implanted sensors. Nevertheless, the FSCV voltage scanning artifacts will still be transmitted to the EPhys recording sensors through the conductive tissue, with the voltage attenuating as a function of distance between the sensors.

Here, we developed a temporal interpolation algorithm to extract neuronal spike activity from EPhys recordings with concurrent FSCV, without the need for additional hardware. We leverage the periodicity of the FSCV scans to detect the timing of these signals and interpolate these artifacts away in the EPhys recording. The algorithm was extended for use to remove 60 Hz line noise along with its harmonics. Removing these artifacts is necessary as they frequently are indistinguishable from physiological spikes and may be falsely identified as such. We characterized the recovery rate of our spike extraction techniques by applying simulated FSCV scan artifacts onto isolated EPhys recordings, demonstrating the ability to extract a large percentage (84.5%) of the original spikes. These methods were further validated in task-performing monkeys, where we were able to capture behaviorally-relevant computations of synchronous dopamine and spike activity from functionally diverse neurons in the striatum. In summary, the techniques described here introduce a new way to decipher the interactive relationship between neuromodulator signaling and surrounding neuronal activity and how these coordinate activities influence and/or are shaped by ongoing behavior.

## Materials and Methods

### Animal procedures

Measurements and experimental methods used in this study were obtained from one female Rhesus monkey (approximately 8.5 years old and weighing approximately 10 kg, at time of recordings). All experimental procedures were approved by the Committee on Animal Care of the Massachusetts Institute of Technology and are described in a previous report (Schwerdt et al., 2020).

### Behavioral task

The monkey performed a visually guided reward-biased task as described previously (Schwerdt et al., 2020). Briefly, each trial consisted of an initial central cue (C) that the animal had to saccade to and fixate on for 1.6 s. A peripheral target (T) cue appeared on the left or right of the screen after the C extinguished. The animal had to saccade and fixate on the T visual cue for 4 s to receive a small or large reward (RW), which consisted of 0.1 – 0.3 mL or 1.5 – 2.8 mL of liquid-food, respectively. The association between target direction (left or right) and reward size (small or large) was switched every 20–30 trials.

### ECP recording setup

Both EPhys and EChem-FSCV measurements were made from CF sensors, fabricated following previously published methods (Schwerdt et al., 2017), implanted in the monkey striatum (**Fig. 1**). The setup is described in a previous publication (Schwerdt et al., 2020) and a brief summary is provided here. CF electrodes consisted of either conventional silica tube-threaded electrodes (silica-CF) or parylene-encapsulated electrodes (py-CF). CF sensors were mounted onto microdrives atop a custom-designed chamber (Gray Matter Research) that was affixed to the monkey’s skull. The sensors were inserted into the brain through guide tubes (Connecticut Hypodermics, 27G XTW) and lowered to the striatum (i.e., caudate nucleus, CN, and putamen). Sensors were individually connected to either standard electrophysiological (EPhys) recording system (Neuralynx, HS-32) or an FSCV system (obtained from S.B. Ng-Evans at University of Washington) as selected on a day-by-day basis. EPhys signals were referenced to multiple tied stainless-steel wires inserted in the tissue above the dura mater (A-M Systems, 790700). Ag/AgCl electrodes were implanted in the epidural tissue and/or in a white matter brain region to serve as the FSCV reference.

Dopamine concentration changes were recorded from implanted sensors using a 4 channel FSCV system (from S. B. Ng-Evans at University of Washington). This consisted of a transimpedance amplifier headstage to convert and amplify electrochemical current to voltage, and a computer to control the applied voltage scans and record and store current. This system generated a triangular voltage ramping from –0.4 V up to 1.3 V and back to –0.4 V, with a scan rate of 400 V/s. This was applied at a sampling frequency of 10 Hz to the connected electrodes. A holding potential of –0.4V was maintained at the electrode between scans. Background-subtracted color plots were generated by plotting the relative current change as color, applied scan voltage on the y-axis (i.e., –0.4 to 1.3 to –0.4 V), and time (i.e., each scan at 100 ms intervals) on the x-axis.

EPhys recording was performed using a standard electrophysiology system (Neuralynx, HS-32). The recording settings were configured as follows: input range of ± 1 mV, sampling rate of 30 kHz, and band pass filter with a passband at 0.1 – 7,500 Hz. This system received the timestamps for behavioral task events as generated from the VCortex behavioral system. A digital messaging system (Neuralynx, NetCom Router) allowed transmitting of trial-start events from the EPhys system to the FSCV system to provide shared timestamps between the FSCV and EPhys systems.

### Dopamine concentration estimation

Dopamine concentration changes ([ΔDA]) were approximated using principal component analysis (PCA), as previously described (Schwerdt et al., 2017). Briefly, each FSCV scan produces a cyclic voltammogram (CV) (i.e., current vs. voltage plot), which is background subtracted to remove the larger current contributions associated mainly with nonfaradaic processes and to distinguish the smaller current changes related to chemical redox. The background subtraction usually occurs at an arbitrary reference time (i.e., alignment event such as the T cue) to provide a uniform reference for all task-modulated signals on each trial. These background-subtracted currents are projected onto the principal components computed from standards of dopamine, pH, and movement artifact as previously established (Schwerdt et al., 2017). CVs that produced an excessive variance (*Q*) above a tolerance level (*Q*_*α*_) could not be accounted as a physiological signal and were nulled automatically (assigned NaN values in MATLAB). Signals were also nulled where CVs were correlated to movement artifact standards (*r* > 0.8). These strict procedures ensure that at least 90% of the signals are identified as dopamine, with a false negative rate of less than 30%, and 0% false positives.

### Spike sorting

Spike sorting was performed manually in commercial software (Plexon, Offline Sorter) following standard protocols (Feingold et al., 2012). Raw EPhys data were first high pass filtered using a Butterworth filter with a cut-off frequency of 250 Hz (4-pole, forward only). A negative threshold was set to identify negative-going crossings of the action potential waveforms. Waveforms were extracted with a length of 1.6 ms (48 samples) using a prethreshold period of 0.267 ms (8 samples). These waveforms were then visualized in terms of their energy, non-linear energy, and their projections onto principal components (PC) space (e.g., PC1 vs PC2). Waveforms were first invalidated based on the prominence of energy and non-linear energy features, which enhances visualization of artifact-like signals such as large transient glitches created by electrostatic discharge as well as from the reward delivery peristaltic pump. Waveforms were then manually invalidated in the PC space if they looked like the artifacts mentioned previously and did not resemble physiological spike signals (e.g., high-frequency changes in voltage). PCs were recalculated after invalidating waveforms to distinguish potential clusters based on the variances calculated for valid waveforms. Clusters were manually drawn on the PC space when a distinct boundary was observed between the projected points. These clustered waveforms were then exported for plotting average waveforms and histograms in MATLAB (Mathworks, 2023b). Interspike interval (ISI) histograms were generated using 1 or 2 ms bin widths to visualize the relative distribution of spike firing intervals, which is used as a parameter to distinguish different cell-types (Aosaki et al., 1994).

### Temporal interpolation methods to extract spike data

We developed a simple automated algorithm to reliably extract extracellular action potentials by performing linear interpolation over the FSCV artifacts embedded in the EPhys waveforms in the time-domain. These methods were implemented in MATLAB (Mathworks, 2023b). The artifact shape and amplitude depends on a number of factors, including the distance between the FSCV recording electrodes and the EPhys electrodes, as well as the characteristics of the EPhys electrodes (e.g., impedance, and other nonlinear electrode-tissue interface properties) (Nag et al., 2015). A flowchart of the algorithm is illustrated in **Fig. 2** and details are as follows. 1) The first step in the algorithm is to get a first approximation of the timings of the artifacts. Signals from multiple EPhys electrodes are averaged to enhance the artifact peaks that are coherent across channels, and to reduce background spike and LFP fluctuations that may inadvertently trigger threshold crossings used in subsequent steps (**Fig. 3A** and **B**). 2) The averaged signal is bandpass filtered (BPF) at 10 – 100 Hz, and then its absolute value is taken.

**Figure 2.**
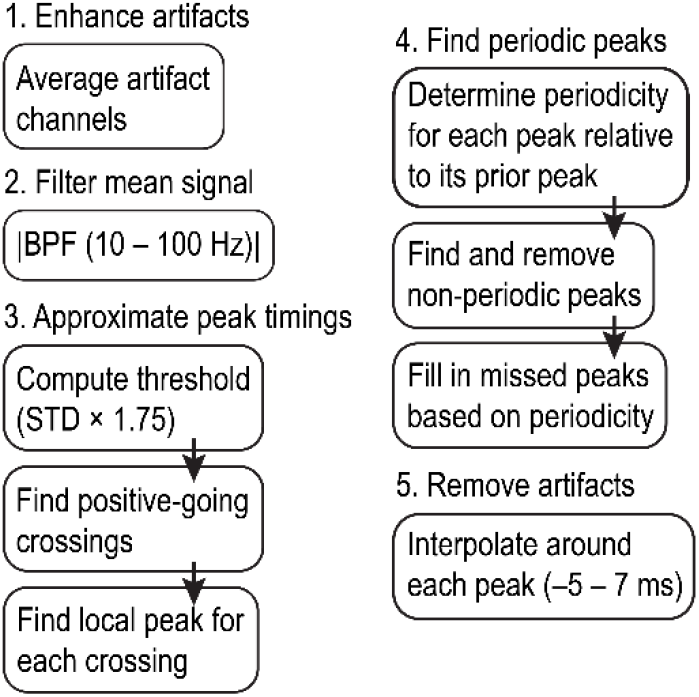
Flow chart of temporal interpolation algorithm used to extract spike activity from ECP recordings. Details may be found in the text.

**Figure 3.**
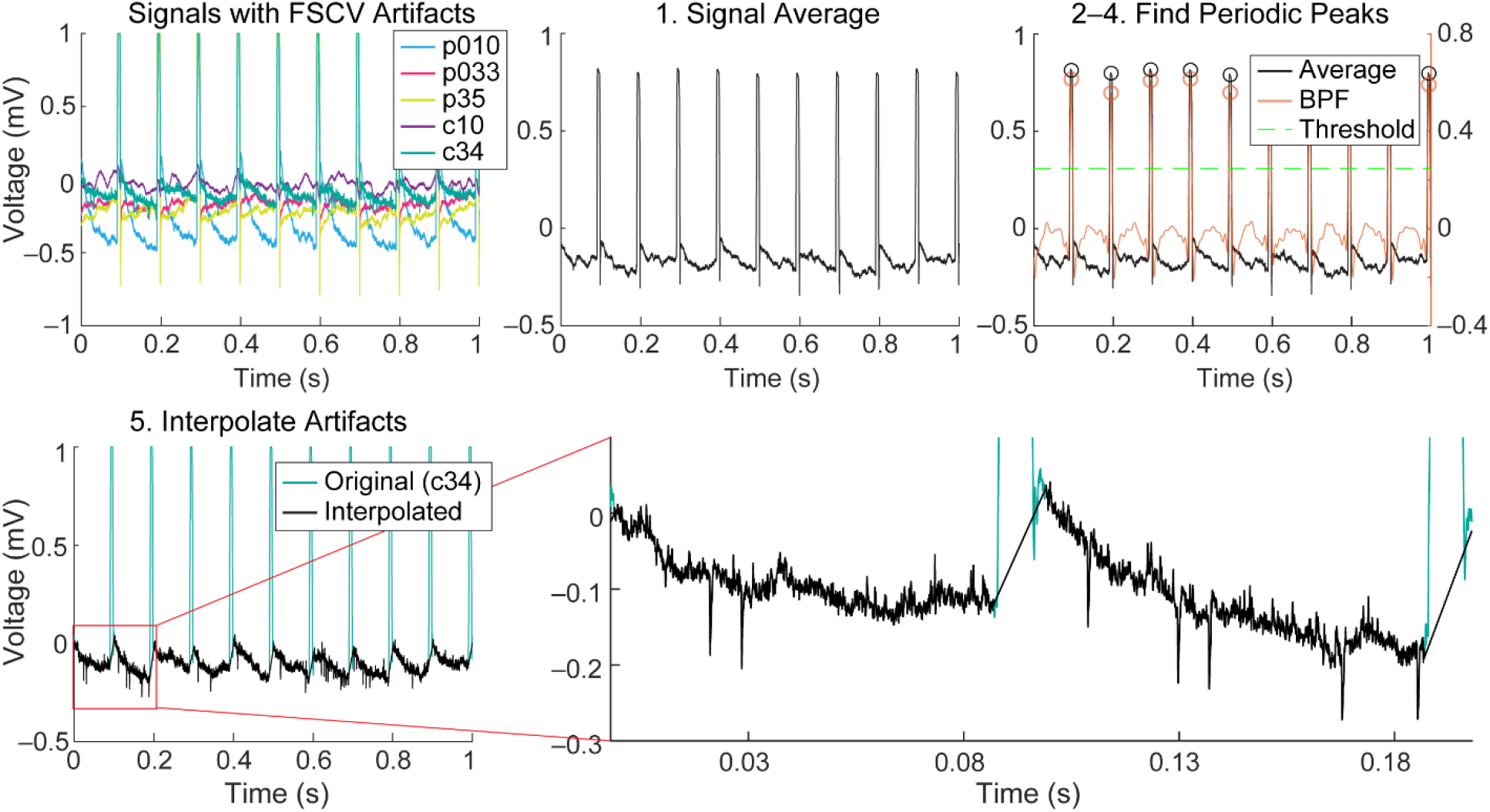
Temporal interpolation process applied to an example ECP recording. Signals from 5 electrodes containing FSCV scan artifacts are plotted in the first panel (top-left) with electrode sites labeled in the legend (c denotes CN site and p denotes putamen site). All these signals are then averaged (step 1 in interpolation algorithm) (top-middle). The averaged signal is bandpass filtered (BPF) and a threshold is computed (1.75 x STD) to find the positive-crossings and local peaks for each of these (circles) (top-right). Only periodic peaks are retained (or added if they did not cross the threshold initially). Linear interpolation is performed around each of these identified peaks using a window of –3 – 7 ms for each electrode channel. An example is shown for site c34 in the bottom-left plot, as well as a close-up on the bottom-right, to visualize recorded unit spike activity.

This procedure further enhances the artifact peaks over surrounding signals. 3) The artifact peak timings are identified by first finding positive-going crossings over a threshold, a multiple (i.e., 1.75) of the standard deviation of the previously calculated signal. The local peak directly following each positive crossing is identified and stored. 4) Only peaks demonstrating periodicity according to the expected FSCV scan frequency (10 Hz) are retained to prevent unwanted removal of physiological signals. This is done by checking the periodicity of each peak relative to its prior peak. Any non-periodic peak is removed from the stored variable containing a list of threshold crossing peaks. Furthermore, missing peaks are added based on the periodicity of the peaks captured before or after the window in which peaks were not detected. This ensures that artifacts that exist below the initial detection threshold are captured and resolved to minimize erroneous physiological data. 5) Finally, linear interpolation is performed on the signal around each identified peak with a window of –5 to 7 ms. This window was empirically determined through incremental (0.5 ms) increases beyond the defined 8.5 ms scan window applied in FSCV. The wider window accounts for recovery time of the amplifier as well as low pass filtering and resulting broadening of the signal through the tissue. This interpolation removes the artifact, preventing it from being detected falsely as a spike during subsequent high pass filtering and thresholding steps used for spike sorting.

This algorithm was further extended to remove artifacts caused by line noise at 60 Hz and its harmonic frequencies (e.g., 120 Hz) (**Extended Data Fig. 3-1**). Such line noise interference was frequently present in our experimental setup, most likely due to the shared ground between EPhys and EChem systems. Removing this interference only required a simple modification to the previous algorithm, where the second filtering step was replaced by a high pass filter (300 Hz cutoff) as this accounted for the higher-frequency content of the noise, and that the threshold computation was performed by taking a multiple (i.e., 8) of the average of the filtered signal rather than its standard deviation.

These interpolation algorithms do not require a clock input to provide timings of the FSCV scans and instead relies on the periodicity of the signal and its appearance on multiple EPhys recording electrodes. The periodicity of the signal is a key input into the algorithm as simple thresholding methods used to remove large glitches or artifacts may inadvertently remove physiological signals. Template methods (Banaie Boroujeni et al., 2020; Hashimoto et al., 2002; Nag et al., 2015; O’Shea and Shenoy, 2018; Wichmann and Devergnas, 2011) frequently used for removing artifacts caused by electrical stimulation are not able to account for the range of variability in the shape of the FSCV artifact, which depends on the distance between the electrodes and the electrode properties. These algorithms are available at github.com (https://github.com/hschwerdt/extractFSCVspikes).

## Results

### Validation of spike extraction methods

Our algorithm was validated by adding synthetic FSCV artifacts onto EPhys recordings made without FSCV (i.e., EPhys-only) (**Fig. 4**). FSCV artifacts were emulated following methods previously established for validating LFP extraction algorithms in similar ECP measurement configurations (Schwerdt et al., 2020). Three different types of artifacts were simulated: resistive (R), resistive-capacitive (RC), and rail (i.e., saturating). These differences reflect characteristics of the artifact duration and amplitude observed in EPhys recordings made during concurrent FSCV, and arise due to differences in distance between electrodes, as well as the electrode properties. Spike sorting was done on the EPhys-only recording, before adding artifacts, and then again after adding artifacts and implementing our interpolation algorithms. Spike recovery rate was defined as the percent of physiological spikes retained after interpolating the artifacts.

**Figure 4.**
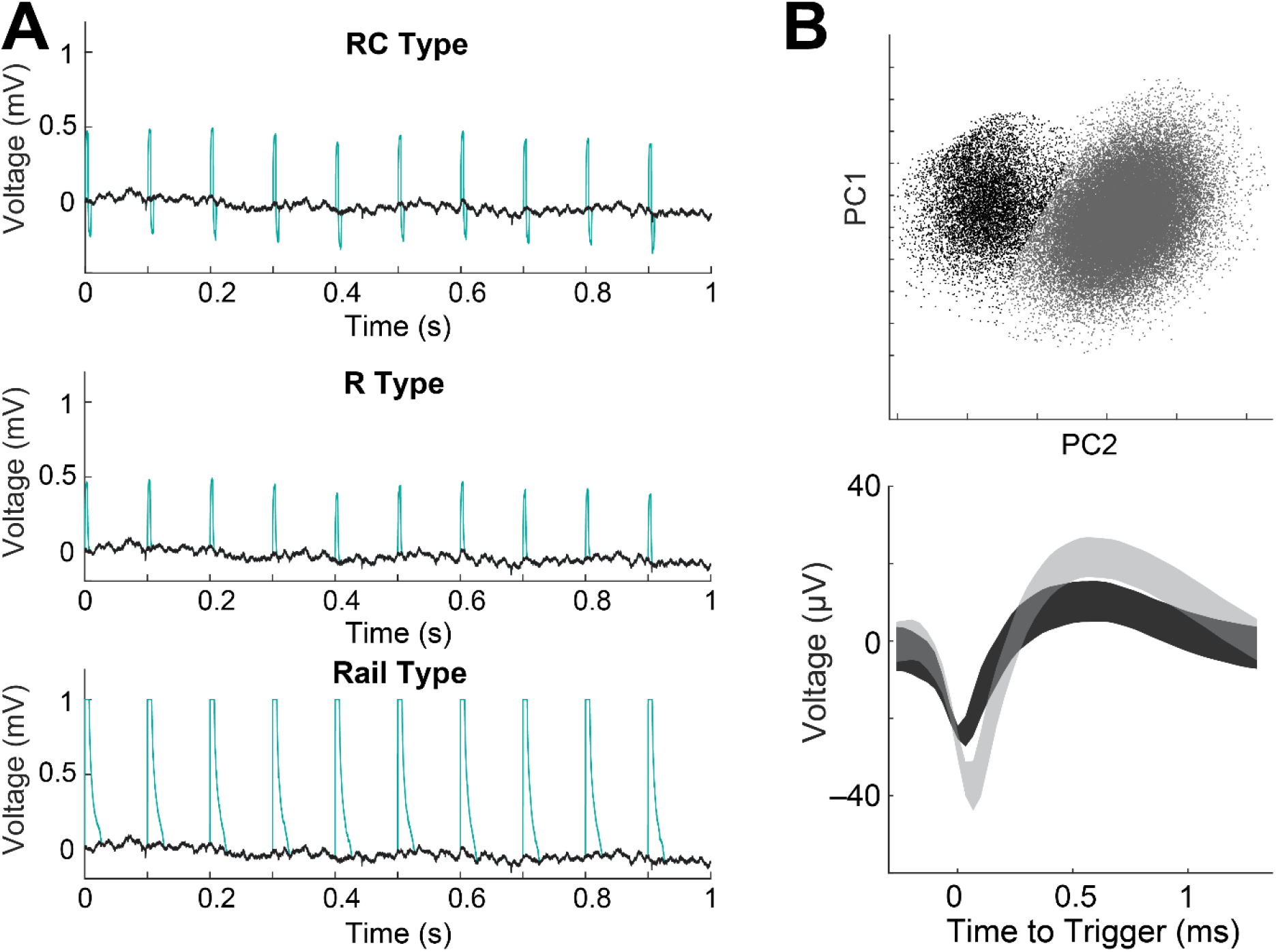
(A) Validation setup showing 3 different simulated FSCV artifact types (RC, R, Rail) (green trace) added onto EPhys-only recording (black trace) (session 65B and site p32). (B) (Top) Spike waveforms projected onto PC space (top) with colored clusters. (Bottom) Average spike waveforms for drawn clusters from the PC space (top). Shading represents +/- SD.

Simulation of each of the three types of artifacts was implemented individually on 5 recorded channels from 3 different sessions and the results are shown in **Table 1**. We calculated the spike recovery rate for each type of artifact being interpolated with our algorithm. We found that the spike recovery rate was 84.5% as averaged for all artifact types. This is equivalent to a loss of 15.5%, which was a reasonable result given that the 8.5 ms FSCV scans make up 8.5% of the recording, and our interpolation occurs over a wider 12 ms and therefore 12% of the recording. The average rates for interpolating each type of artifact were 88%, 85.8%, and 79.6% for the R, RC, and rail type artifacts. Spike recovery rates were the worst for interpolating rail-type artifacts as expected given their broader width that encompasses the amplifier recovery time after its input saturates. This is important to consider when designing higher-density configurations of CF sensor arrays as minimizing the distance between sensors will increase the artifact amplitudes. Code and data used for simulating artifacts and validation are available at zenodo (10.5281/zenodo.10396372).

**Table 1.** Validation results showing percent recovery of spikes extracted after interpolating simulated artifacts relative to spikes extracted from clean EPhys recording for the different types of artifacts (R, RC, and Rail).

| Session | Channel | R recovery | RC recovery | Rail recovery |
| --- | --- | --- | --- | --- |
| 65B | p23 | 87.13 | 86.39 | 79.55 |
|  | p32 | 87.23 | 86.9 | 83.51 |
| 109B | p35 | 88.66 | 86.07 | 76.07 |
|  | c34 | 91.78 | 84.48 | 81.02 |
| 161b | p36 | 85.35 | 85.06 | 77.94 |

### Measurements of cell selective spike activity

Extracted spike activity was analyzed as measured from implanted CF sensors in the CN and putamen during concurrent FSCV in a task-performing monkey (**Figs. 5** and **6**). Striatal units have been shown to display specific waveform shapes and firing characteristics depending on the cell type classification (Apicella, 2002; Hikosaka et al., 1989; Thorn et al., 2010; Yamada et al., 2004). Our units displayed distinct characteristics that largely resembled either putative medium spiny neurons (MSNs) or tonically active neurons (TANs), as classified in prior work. TANs display a longer after-hyperpolarization (i.e., broader shape) (**Fig. 6E**) in comparison to MSNs (**Fig. 5B**) (Apicella, 2002; Hikosaka et al., 1989). Furthermore, MSNs are known to fire scarcely (< 1 Hz) and increase sharply (i.e., burst) in response to relevant behavioral events, which is noticed in the peak spike counts amongst the low interspike intervals (ISIs) in the plotted ISI histograms (**Fig. 5B** and **Fig. 6B**). On the other hand, TANs maintain spontaneous firing rates of 2 – 12 Hz and show transient pauses in response to a variety of stimuli or events (Aosaki et al., 1995, 1994; Apicella, 2002; Hikosaka et al., 1989). TANs may also display up to two peaks in their ISI histograms (Aosaki et al., 1994), which was also observed in our unit (**Fig. 6E**). These cell type distinctions may also be perceived in the spike rate histograms plotted in association with behavioral events (**Fig. 5C** and **Fig. 6A** and **D**), where the average spontaneous firing rates outside of trial bounds (i.e., before the central start cue and after the outcome) are higher for putative TANs (**Fig. 6D**) than for MSNs (**Fig. 5C** and **Fig. 6A**). The event-related discharges and pauses of putative MSNs and TANs, respectively, may be observed in these plots. These results collectively demonstrate our ability to resolve cell-type specific spike activity as measured during concurrent FSCV electrochemical recording.

**Figure 5.**
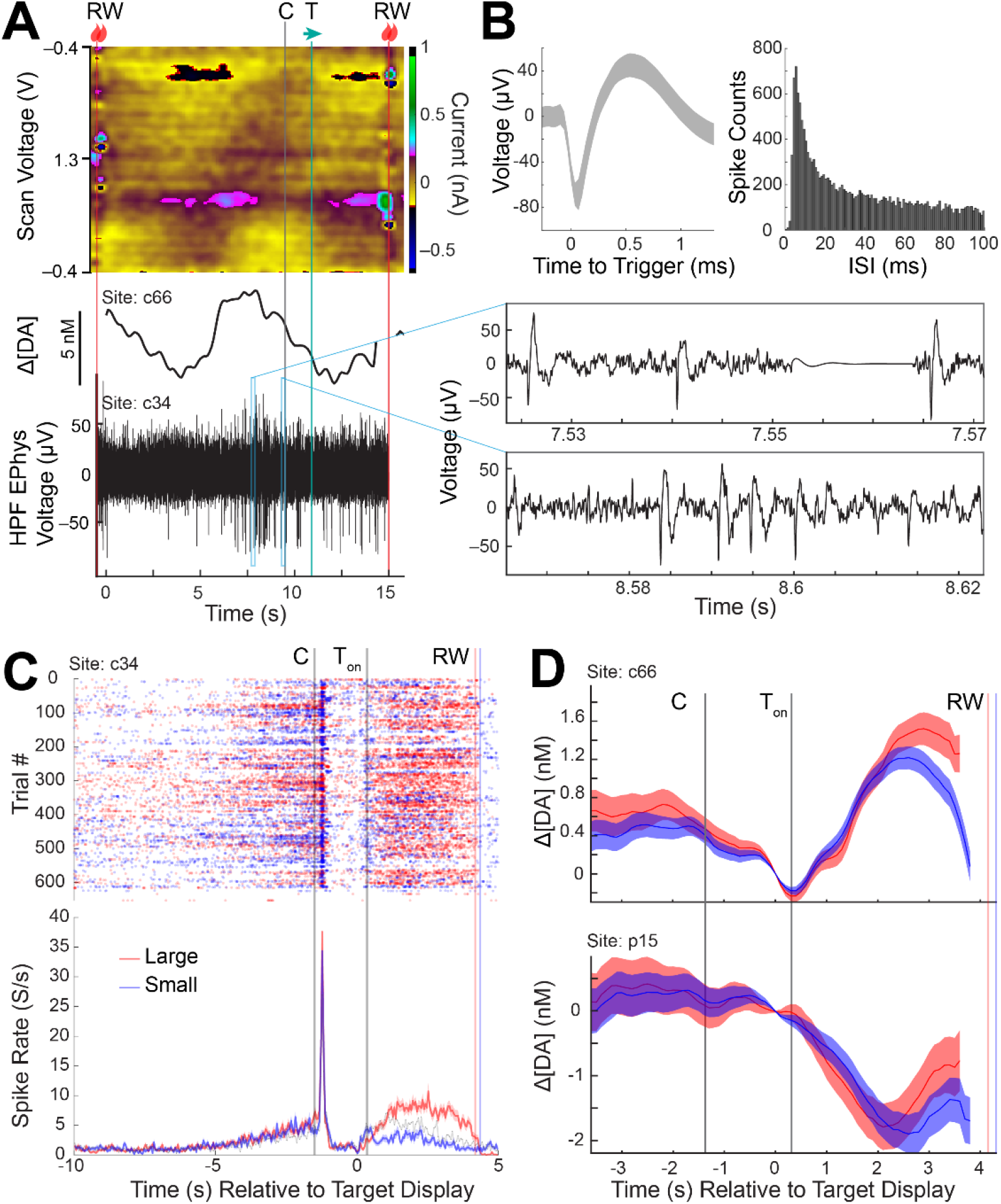
Analysis of task modulated signaling from concurrent recordings of dopamine and spike activity from three sites in the CN and putamen measured from a single session (session 127). (A) (Top) FSCV color plot, (middle) PCA-computed dopamine concentration change ([ΔDA]), and (bottom) concurrent measurements of electrical neural activity high pass filtered (HPF) to visualize spike action potentials. Two windows (blue rectangles) are magnified to show the individual spike action potential waveforms (right). Task events are labeled following notation in **Fig. 1**. (B) (Left) Average waveform of unit detected (shading represents +/- SD). (Right) ISI histogram of the detected unit. (C) (Top) Trial by trial raster plot of spike activity (dot) measured in the CN (c34) as aligned to the peripheral target display event (0 s). T_on_ represents the average time from the peripheral target display event at which the monkey begins fixation on the peripheral target. (Bottom) Average spike rate for large and small reward trial conditions (shading represents +/- SE). Large and small reward trials are denoted by red and blue colors, respectively. (D) Dopamine concentration changes measured at neighboring sites in the CN (c66) and putamen (p15) as aligned to the same events as (C) for large and small reward conditions (shading represents +/- SE). Color coding is the same as in (C).

**Figure 6.**
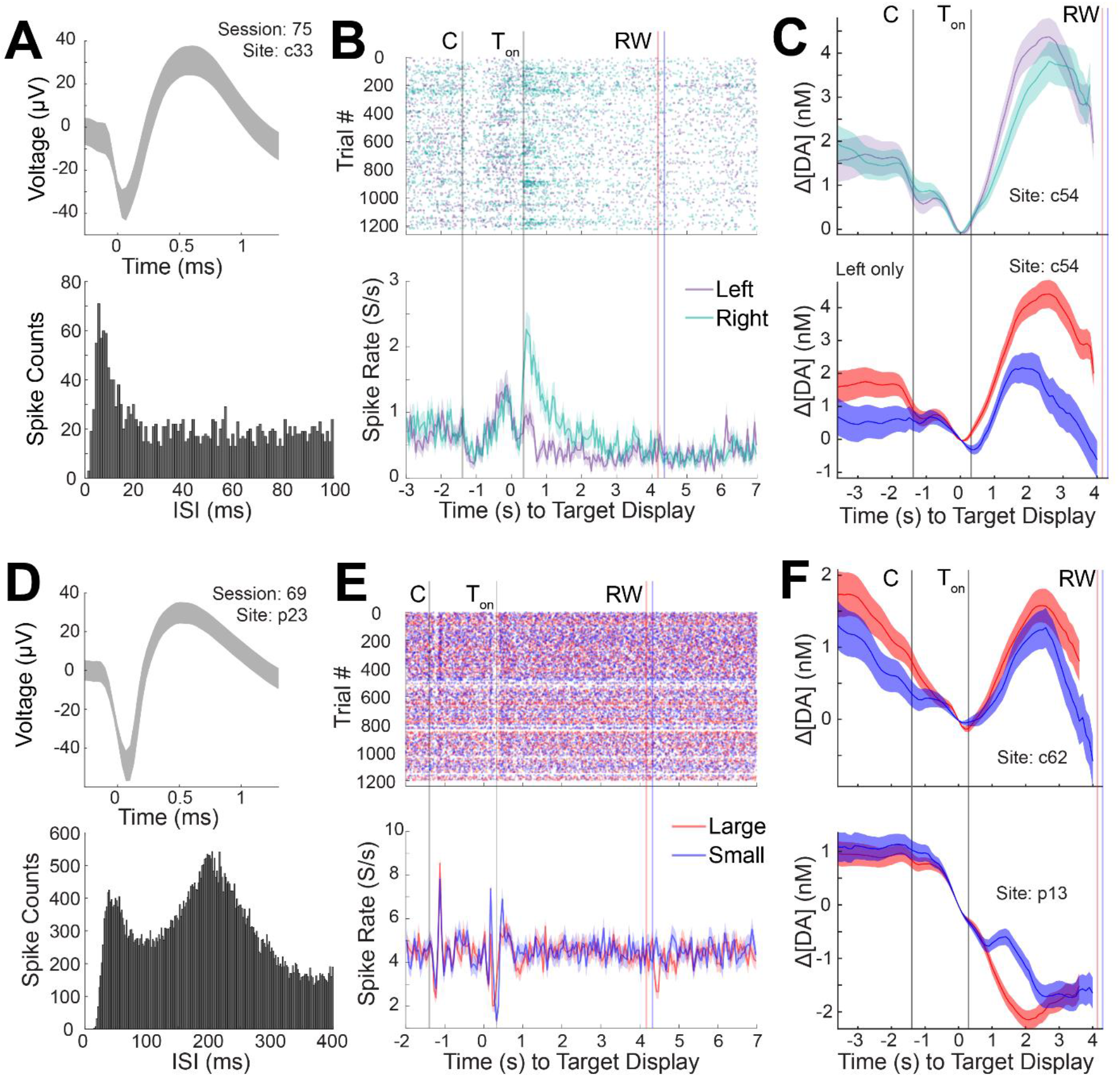
Dopamine and spike activity recorded concurrently from a single session show diverse responses to behavioral events related to reward size and spatial target direction. (A) Average waveform of a putative MSN (top) and its ISI histogram (bottom) as recorded in the CN (c33). (B) Raster plot of spike activity for the unit in (A) as plotted in **Fig. 5C**, except for left (purple) and right (green) peripheral target conditions, demonstrating higher neural responses to gaze of the right peripheral targets than to left targets. (C) Dopamine concentration changes measured at a neighboring site in the CN (c54) displaying oppositive sensitivity to target direction (higher for left than for right target) in comparison to the unit in (B) (top), and stronger modulation by reward size (bottom). Color coding is the same as in (B) and **Fig. 5C**. (D) Same as (A) except for another unit (putative TAN) recorded in a separate session (69) and site in the putamen (p23). (E) Same as (B) except for large (red) and small (blue) reward conditions. No distinction is observed in the cell firing for the reward size or target direction. (F) Dopamine concentration changes measured at neighboring sites in the CN (c62) and putamen (p13) where stronger modulation by reward size is observed in comparison to the unit response in (E). Color coding is the same as in (E).

### Behaviorally relevant measurements of dopamine and spike co-activity

We analyzed how synchronously recorded dopamine and spike activity were modulated by behavioral task events related to reward size, eye movement direction, and visual cues. Multi-modal measurements were made in a monkey performing a task where eye movements were made to left or right targets to receive liquid-food rewards. The size of the reward (i.e., large or small) depended on the target location (i.e., left or right) and this was switched every block to counterbalance reward size and movement direction variables. More details of the task are described above in “Behavioral task”.

Spike activity of a putative MSN recorded in the CN (c34) was shown to increase significantly in response to the appearance of the initial central start cue. This neuron also showed significant modulation by the size (large or small) of the upcoming reward (**Fig. 5C**). Such reward-size modulation is similar to that observed in prior experiments (Cromwell and Schultz, 2003; Kawagoe et al., 2004). These reward-size activities were maintained for a sustained period throughout the period from the target cue onset to the reward delivery, possibly reflecting ongoing functions related to invigoration (Cromwell and Schultz, 2003). Furthermore, pre-cue anticipatory activity is observed in this neuron, also seen in previous work (Kawagoe et al., 2004), and has been linked to facilitating subsequent movements. We found that dopamine was also modulated by reward size anticipation in a neighboring site (c66) in the CN, but not in our putamen site (p15) (**Fig. 5D**). Dopamine signals were not modulated by the initial central cue in either site.

In another CN site (c33) and session, a putative MSN’s spike activity was found to be target direction sensitive, displaying higher firing rates during gazes to right (ipsilateral) targets in comparison to left (**Fig. 6A**). Unlike the previous unit, this neuron was not modulated by reward size. Such spatial selectivity has been observed of striatal units recorded in similar eye movement tasks (Kawagoe et al., 2004). Synchronously recorded dopamine signals, on the other hand, were modulated by both reward size and target direction (**Fig. 6C**). However, dopamine was slightly higher for contralateral targets.

A putative TAN recorded in the putamen (p23), also in a separate session, displayed transient decreases in spike firing in response to displayed visual cues (i.e., initial central cue and peripheral target) followed by a rebound increase, replicating event-related pause activity frequently observed in TANs (**Fig. 6D**) (Aosaki et al., 1995, 1994; Apicella, 2002; Hikosaka et al., 1989). The neuron did not discriminate between reward size or movement direction. These patterns of observations are in line with prior work (Apicella, 2002; Hikosaka et al., 1989; Thorn et al., 2010; Yamada et al., 2004), which have largely attributed these signals to representing the motivational value or arousing aspects of external stimuli. Striatal dopamine in the putamen and CN was modulated by reward and target cue direction (**Fig. 6F**), similar to our previous examples and other reports (Schwerdt et al., 2020).

## Discussion

Methods for recording and analyzing extracellular action potentials as measured concomitantly with FSCV-based dopamine signals were developed and validated in this study. Minimal additional hardware was required beyond standard EPhys and FSCV instrumentation to develop our ECP system. Spike extraction was carried out offline using simple custom-made temporal interpolation algorithms. These algorithms leveraged the periodicity of the FSCV voltage scans to remove the interfering FSCV voltage scan artifacts off the EPhys recordings. We validated the extraction technique using artificial artifacts to ensure that spikes were retained with high fidelity. ECP recording was performed on CF sensors implanted in the striatum of a monkey to uncover the co-active dopamine and spike signals underlying reward and movement behaviors. We further demonstrated the ability to distinguish different neuronal cell-types as well as behavioral functions of our recorded units.

Spike recovery rates of 84.5% were achieved using the temporal interpolation and spike extraction techniques developed in this study. This high recovery rate allowed us to distinguish behavioral correlates of multiple identified neural units, and to differentiate dopamine and neuronal functions. We found that unit activity and dopamine displayed very different patterns of signaling in response to behavioral events in our task. Striatal neurons are known to represent a multitude of parameters related to conflict decision-making (Amemori et al., 2020), learning (Desrochers et al., 2015), and other behavioral functions (Apicella, 2002; Hikosaka et al., 1989; Thorn et al., 2010; Yamada et al., 2004) (Aosaki et al., 1995, 1994; Apicella, 2002; Hikosaka et al., 1989). On the other hand, dopamine has been largely attributed to a role in reward valuation and prediction error signaling (Schultz et al., 1997). Only recently has dopamine been shown to also provide a prolific representation of motor and sensory variables, outside of simple reward variables (Coddington et al., 2023; Menegas et al., 2018; Schwerdt et al., 2020). A remaining question is how dopamine influences the activity of nearby neurons (Sippy and Tritsch, 2023), such as in the form of plasticity (Brzosko et al., 2019; Shindou et al., 2019), and how these interactions modify or are modified by behavior. The reverse of this, understanding how neuronal activity influences dopamine release (Threlfell et al., 2012), also remains an unresolved question that may be addressed through multi-modal measurements such as those demonstrated in this work. Nevertheless, standard trial-averaged computations, as used here, may provide limited insight of such potential interactions. Furthermore, measurements of dopamine and neuronal activity should be performed at local sites given dopamine’s spatially heterogenous operations (Hamid et al., 2021; Schwerdt et al., 2020). Higher recovery rates may be possible for non-saturating artifacts (i.e., R and RC type) using techniques that may potentially be applied to subtract out the artifact and recover spikes during the artifact (Banaie Boroujeni et al., 2020; Hashimoto et al., 2002; Nag et al., 2015; O’Shea and Shenoy, 2018; Wichmann and Devergnas, 2011).

One of the limitations of our current configuration is that dopamine and neuronal signals are measured from separate, and distal (> 1 mm), sensors. Single-sensor systems have been previously developed to combine FSCV transimpedance and EPhys voltage amplifiers with active switching circuitry to measure chemical and voltage signals from the same sensor and site.(Takmakov et al., 2011). These have been applied in rodents to successfully measure electrical-stimulation evoked dopamine and spike activity (Cheer et al., 2005). Nevertheless, as described in the “Introduction”, a single-sensor configuration prevents a negative hold potential from being applied in between applied voltage scans, significantly restricting the sensitivity to measure dopamine and other positively charged molecules (Bath et al., 2000). As far as we are aware, measurements of endogenous (i.e., not electrically stimulated) neurochemical signaling with this single-sensor configuration have not been demonstrated. On the other hand, separating the sensors for FSCV and EPhys enabled concurrent recording of naturally-occurring dopamine and voltage fluctuations during behavior in rodents (Parent et al., 2017) and monkeys (Schwerdt et al., 2020). This decoupled ECP configuration was therefore used in this work.

So far, two variations of a decoupled ECP system have been reported, as far as we are aware (Parent et al., 2017; Schwerdt et al., 2020). The first system attempted to address or reduce the effects of the FSCV artifacts through several hardware and software modifications. This would allow successful measurements of stimulation and pharmacologically evoked dopamine signals in awake and mobile rodents. A relay circuit was used to isolate FSCV scan voltages from the EPhys recording system. In principle, this would be helpful to prevent large voltages from saturating the EPhys input amplifiers when the electrodes are close to each other. Nevertheless, the relay induced larger artifacts than FSCV scan voltages. A timing signal for the scan voltages was sent to both FSCV and EPhys systems, which allowed readily extracting EPhys recorded signals during the window in between the known scan periods, or interpolating away these periods. An additional window of interpolation (4.5 ms) was added around the scan period, similar to the current work, most likely also to remove the effects of capacitive discharge and amplifier recovery time. Furthermore, a lower scan frequency (5 Hz) was used to produce a wider artifact-free window in which lower frequency LFPs could be extracted. However, this limits the temporal resolution of the dopamine measurements. Similar to our current work, the second system utilized the standard 10 Hz scan frequency to maintain a higher temporal resolution to capture the fast millisecond dynamics of dopamine release and clearance, and did not require any clocked input for transmitting timings of the FSCV scans, or other specialized hardware (i.e., relay) (Schwerdt et al., 2020). EPhys and FSCV recordings were synchronized through shared behavioral event codes. This system applied a custom-made algorithm to interpolate away these artifacts in the frequency domain, allowing reliable extrapolation of a broad frequency range (0.1 Hz – 1 kHz) of LFPs with a high correlation to original waveforms (R ∼ 0.99), based on simulated artifacts. However, these algorithms were not capable of reliably extracting extracellular action potentials (i.e., spikes or units) due to the higher frequency content of these signals (up to ∼8 kHz) (Fee et al., 1996). Thus, in this work, time-domain interpolation is used instead of the frequency-domain to preserve the broad frequency content of the spike waveforms, which is imperative to identify individual units via standard spike-sorting methods.

Another potential advantage of our ECP system is the ability to record both unit activity and dopamine signals from the same CF sensor, albeit at different times. This would allow comparisons between spike and dopamine activity from the same site, as measured from different recording sessions. This could be useful for making associations between dopamine and neuronal activity during well-maintained and reproducible behaviors across sessions. Nevertheless, ideally, such measurements would occur at the same time in order to infer temporal correlations between recorded multi-modal signals. Measurements from juxtaposed electrode contacts may provide the needed focal site-specific metrics of interacting signals.

In summary, we developed a simple system combining standard EPhys and FSCV instrumentation for synchronous measurements of neuronal spike and dopamine signaling for use in nonhuman primates. The system could easily be adopted for use in other species, such as rodents (Parent et al., 2017) and humans (Kishida et al., 2016). Combinatorial methods such as those described here are needed in behavioral experiments to help resolve important questions related to the physiological mechanisms of plasticity and the interactions between neuromodulators and target neurons that regulate ongoing behavior.

## Data Availability

All source data are available at zenodo.org (**10.5281/zenodo.10397754, 10.5281/zenodo.10397773**).

## Code Availability

MATLAB code used to analyze data are available at GitHub (https://github.com/hschwerdt/extractFSCVspikes, https://github.com/hschwerdt/fscvartifactcreation).

## Acknowledgment

This work was supported by NIH NINDS (R00 NS107639 to H.N.S.), the Michael J. Fox Foundation for Parkinson’s Research (MJFF) and the Aligning Science Across Parkinson’s (ASAP) initiative (ASAP-020-519 to H.N.S.), NIH/NIMH (P50 MH119467 to A.M.G), the Army Research Office (W911NF-16-1-0474 to A.M.G.), and Mr. Robert Buxton (to A.M.G.). MJFF administers the ASAP-020-519 on behalf of ASAP and itself.

## Author Contributions

H.N.S. and U.A. designed and validated methods. H.N.S. performed *in vivo* experiments. H.N.S, U.A., J.C., and R.M. analyzed data. H.N.S., D.J.G., and A.M.G. guided methods and experiments. H.N.S., U.A., and J.C. wrote manuscript with comments from all other authors.

## Figures and Captions

**Extended Data Figure 3-1.**
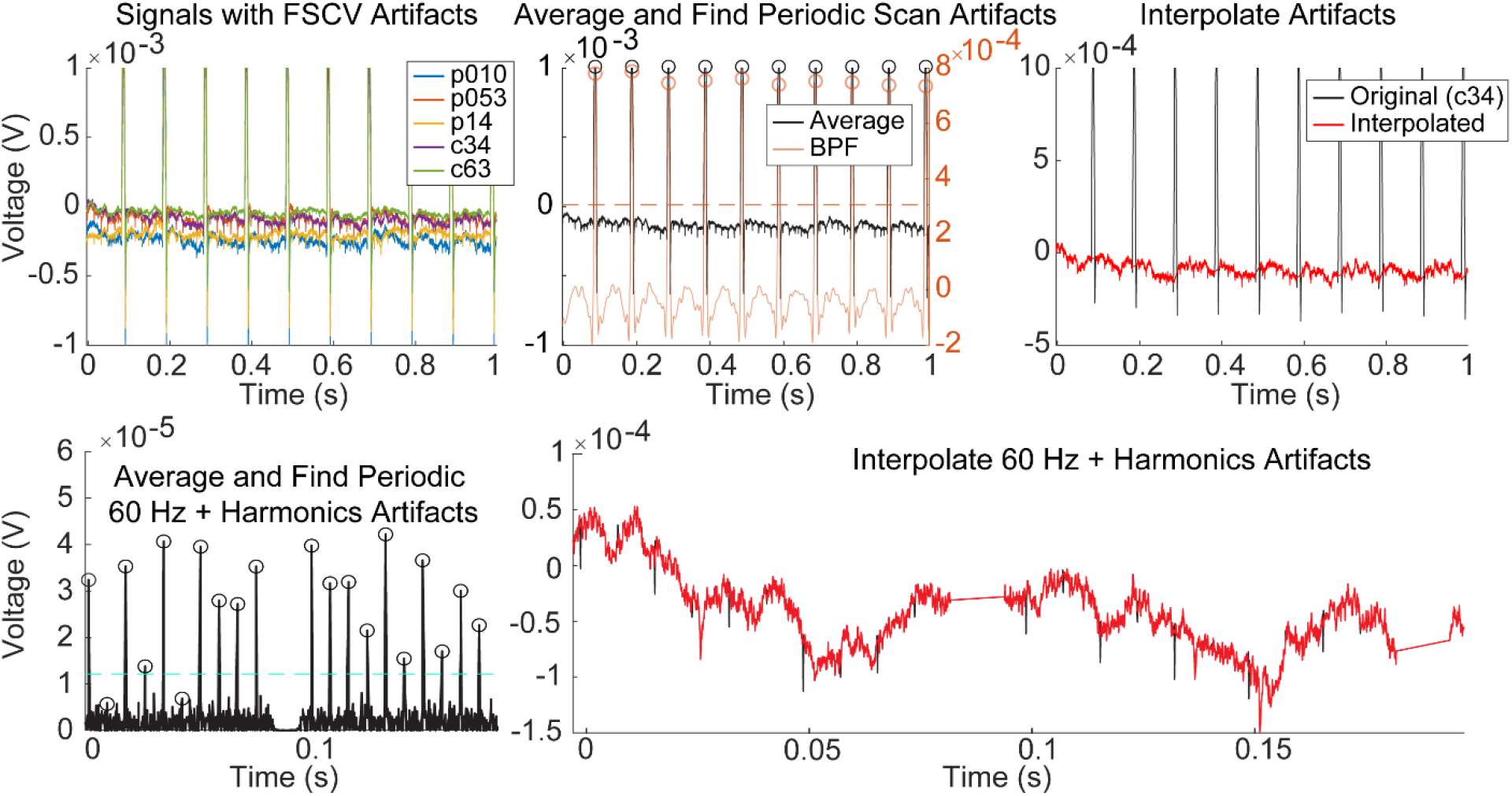
Same as **Fig. 3**, but demonstrating an example with 60 Hz + harmonics noise and the additional steps to remove these signals. Bottom left plot shows the noise as enhanced after high pass filtering. Bottom right plot shows the signal after interpolating both 60 Hz harmonics noise and the FSCV artifacts.

